# A General Biphasic-Bodyweight Model for Scaling Basal Metabolic Rate, Glomerular Filtration Rate and Drug Clearance from Birth to Adulthood

**DOI:** 10.1101/2021.07.25.453668

**Authors:** Teh-Min Hu

## Abstract

Understanding the maturation process of human physiology and metabolism has broad medical and pharmaceutical implications. Age and bodyweight are frequently considered as separate variables in modeling the dynamical changes of human organ functions and of drug clearance from birth to adulthood. The objective of this study is to propose a unified, continuous and bodyweight-only equation to quantify the changes of human basal metabolic rate (*BMR*), glomerular filtration rate (*GFR*) and drug clearance (*CL*) from infancy to adulthood. The *BMR* datasets were retrieved from a comprehensive historical database of male and female subjects (0.02 to 64 years). The *CL* datasets for 17 drugs and the *GFR* dataset were generated from age-incorporated maturation-and-growth models with reported parameter values. The model used in the simulation is independent of the proposed model. A statistical approach was used to simulate the model generated *CL* and *GFR* data for a hypothetical population with 26 age groups (ranging from 0 to 20 years). Besides, individual *CL* data for one drug, and sparse *PBPK*-modeled *CL* values for two drugs were also included for analysis. A 4-parameter, mixed-allometry equation with two power-law functions of bodyweight was proposed and evaluated as a general model using nonlinear regression and dimensionless analysis. All datasets universally reveal biphasic curves with two distinct linear segments on log-log plots. Compared with simple allometry, the biphasic model fits satisfactorily to all datasets (based on Akaike’s Information Criterion and residual plots). The biphasic equation consists of two reciprocal allometric terms that asymptotically determine the overall curvature. The fitting results show a superlinear scaling phase (slope >1; ca. 1.5 – 3.5) below the characteristic bodyweight at the phase transition; and above which, a sublinear scaling phase (slope <1; ca. 0.5 – 0.7) is evident. The phase-transition bodyweight is ranging from 5 to 20 kg (corresponding to 0.5 – 9 years) and the mean value is around 10 kg (∼2 years) for all data sets. The dimensionless analysis generalizes, and offers quantitative realization of, the maturation and growth process. In conclusion, the proposed mixed-allometry equation is a generic model that quantitatively describes the phase transition occurring in the human maturation process of *BMR, GFR* and drug *CL*.

## 1. Introduction

Human metabolic processes undergo continuous changes during the growth and maturation stages (1, 2). It is generally accepted that human physiological and metabolic parameters (*Y*) do not scale linearly with bodyweight (*W*), but scale with a power-law function, in the form of *Y* = *aW*^*b*^, where *a* and *b* are constants). This simple allometric equation is linear in a double logarithmic plot (i.e. log *Y* vs log *W*), which has been used to describe datasets of basal metabolic rate (*BMR*), glomerular filtration rate (*GFR*) and drug clearance (*CL*) (3, 4). However, if data from neonates and adults were pooled for analysis, the resulting plots are usually curved with a steeper early phase, followed by a shallower later phase (3, 5-7). Accordingly, simple allometry with a fixed exponent, or with a specific exponent obtained from the adult data, would systematically overestimate *GFR* or *CL* values among the youngest (3, 7, 8). In the biology field, extensive studies have shown that the metabolic pace of animals or plants shifts at certain points of organism’s life; such shifting exhibits multiple linear phases connected at one or more transitional points in log-log size-scaling plots (9-13). The metabolic shifting phenomenon has long been recognized in human (5), and the quest for a general quantitative model has never ceased (8, 14-17). Thus, the current study addresses the feasibility of using one unified, empirical equation, where bodyweight is the only independent variable, to quantitatively describe the maturation of human physiological and metabolic functions.

The basal metabolic rate (*BMR*), i.e. the rate of energy transformation within the body at rest, is perhaps the most fundamental biological rate. *BMR* is considered as a universal ‘pacemaker’ for biological processes (11). Therefore, allometric scaling of *BMR* has gained much attention and been hotly debated for over a century. In biology, the discussion has been shifted from whether there is a universal scaling law to how intra- and interspecific scaling vary beyond the ‘3/4-power’ law (10). In medicine, human energy requirements are important considerations in clinical and public-health nutrition (4, 18-22). Thus, the WHO issues sex- and age-specific recommendations on human energy requirements (23). The WHO recommendations consist, for each sex, of 3 bodyweight-based linear equations for estimating *BMR*. The equations have different constant values for separate age groups (0-3 yr, 4-10 yr and 11-18 yr), suggesting that *BMR* varies with bodyweight in a nonlinear manner during the growth period. To date, the inherent nonlinearity in human metabolic scaling remains to be addressed by a unified quantitative model.

Glomerular filtration rate (*GFR*) is an important parameter of renal function. In pharmacology and toxicology, *GFR* determines the renal elimination capacity for xenobiotics and environmental toxicants. The information of *GFR* is particularly useful for dosing renally cleared drugs, especially for pediatric patients, because dosing information for children is often limited by ethical and clinical considerations in clinical trials. Accordingly, attempts have been made to predict pediatric *GFR* values from adult’s value using quantitative models (7, 24). Existing data show that *GFR* undergoes continuous change after birth and reaches adult’s level in the early childhood (7, 24). Several models have been proposed to model the growth and maturation of *GFR*, with a focus on predicting *GFR* for neonates or children from adults (6, 7, 24). The previous models have different levels of complexity. The simplest model has the form of simple allometry, where *GFR* scales with bodyweight in a single power-law function, with either fixed or variable exponents (6). More complicated models include both age and bodyweight as the input variables (7, 24). Studies have shown that all models encounter the same uncertainty problem in prospective prediction (6).

Drug clearance (*CL*) is an essential pharmacokinetic parameter that offers direct estimation of drug dose. The acquisition of accurate *CL* information is critical in drug development and in drug therapy. However, the pediatric *CL* data are less attainable, because of the limitation of conducting clinical trials in young subjects. Thus, various prediction models have been proposed to draw quantitative relationship between *CL* and age and/or bodyweight (3, 8, 25-27). The magnitude of *CL* is correlated with body size and determined by the function of drug elimination organs (i.e. the kidneys and liver). Therefore, *CL* values are subjected to changes of renal and hepatic functions during maturation and growth. The *CL* values of many drugs have been compiled from both population pharmacokinetic studies and physiologically-based pharmacokinetic (PBPK) modeling (8, 28-32). Besides, several growth and maturation models have been evaluated for their predictive performance (17, 30, 33-35). Overall, the previous findings are highlighted as follows: First, *CL* scales with bodyweight raised to a power that is variable with age or body weight (16, 28, 30). Second, more profound changes were observed in earlier life than in later life (e.g. neonates to toddlers vs. toddlers to adolescents or adults) (28, 30). Finally, a quantitative birth-to-adulthood model usually includes both age and bodyweight as the independent variables (7, 8, 15, 31, 36, 37), although, for some drugs and in particular age groups, e.g., in younger aged children, clearance can be scaled with bodyweight alone (3, 8, 30, 38-40).

Taken together, human *BMR, GFR* and *CL* values seemed to share similar rate-switching features during the human maturation and growth processes. The aim of this study is to evaluate a continuous, single, bodyweight-only equation for its potential as a unified empirical model for scaling human *BMR, GFR* and *CL* from neonates to adults. The equation is in the form of reciprocal sum of two power-law functions of bodyweight with 4 parameters (so-called mixed-allometry or mixed-power-law equation). All the datasets evaluated in this study were either directly retrieved from the literature or generated from existing age- and bodyweight equations with published parameters. The datasets thus represent the tendency of each specific dependent variable of interest (*BMR, GFR* or *CL*), and the overall goal is not to predict the unknown, but to infer common quantitative understanding from analysis of diverse datasets.

## 2. Methods

### 2.1 *BMR* datasets

The basal metabolic rate (*BMR*) data, in kcal/day, for males and females from birth to adulthood were obtained from the historical database compiled by Schofield (41), The *BMR* data in MJ/day was converted to kcal/day. The database included *BMR* data for 4811 males (0.02 – 52.3 yr, 2.7 – 108.9 kg) and 2364 females (0.14 – 64 yr, 2.0 – 96.4 kg). The data are listed according to divided bodyweight groups (separated by 1 kg up to ∼100 kg). The majority of groups contain >5 subjects (up to 189 subjects).

### 2.2 *GFR* dataset

The glomerular filtration rate (*GFR*) data, in mL/min, were generated using the general equation proposed by Anderson et al. (24) for hypothetical human subjects.

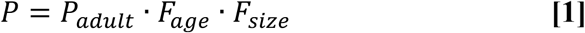

Where here *P* is *GFR* and *P*_*adult*_ is the adult *GFR* value (136 mL/min (7)); *F*_*age*_ is the age function (maturation function), and *F*_*size*_ is the body size function (allometric function):

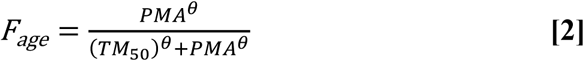

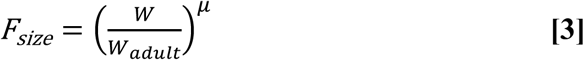

The variables and parameters are defined as: *PMA*, postmenstrual age (weeks); *θ*, the Hill coefficient; *TM*_50_, maturation half-life; *W*, bodyweight of the individual; *W*_*adult*_, adult bodyweight (70 kg); μ, allometric, bodyweight exponent.

The *GFR* dataset was generated for a hypothetical population from 0 to 20 years of postnatal age. The age-bodyweight information was mainly obtained from the Annex 2 in ref. (42). Additional age-bodyweight data for infants <0.25 years were retrieved from ref. (7). A total of 26 age groups were included for the simulation: 0, 0.083, 0.167, 0.25, 0.5, 0.75, 1, 1.5, 2, 3, 4, 5, 6, 7, 8, 9, 10, 11, 12, 13, 14, 15, 16, 17, 18 and 20 years. The mean bodyweight for each age group was either directly retrieved from the reported mean values (7) or estimated from the reported median values (42). Assuming lognormal distribution (43, 44), the median (μ*) was transformed to mean (*μ*) using the following formula (45): *μ* = ln*μ*∗. The standard deviation (*σ*) of lognormal distribution was estimated as: 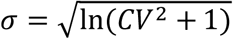, where *CV* is the arithmetic coefficient of variation (46). For each age group, the bodyweight was randomly generated using the age-specific *μ* and *σ*, assuming *CV* = 0.15 in a population (42). This sampling process was conducted using the built-in functions, *RandomVariate* and *LogNormalDistribution*, in Mathematica^®^. The postmenstrual ages (*PMA*) were calculated by adding a gestational age of 40 weeks (24) to the postnatal ages. The resulting (*PMA, W*) data were plugged into Equations 1-3 to generate expected *GFR* values, using the reported parameter values (24): *θ* = 3.4, *TM*_50_ = 47.7 weeks, μ = 0.75. To add variability to the *GFR* values, the lognormal distribution was assumed with *CV* = 0.2. For each age group, *GFR* values for 20 randomly selected subjects were simulated. The data for 26 age groups (a total of 520 data points) were pooled as the *GFR* dataset

### 2.3 Drug *CL* datasets

The first *CL* datasets were retrieved from Björkman’s work (29) on physiologically based pharmacokinetic (*PBPK*) modeling for two drugs, theophylline and midazolam. The *CL* and bodyweight values of both sexes are directly available for neonates, children ages 0.5, 1, 2, 5, 10 and 15 years, and adults. The total *CL* values were estimated from *PBPK*-modeled hepatic *CL*, and/or renal *CL* (if applicable). These *CL* datasets were evaluated in the initial modeling trial for drug *CL*.

Next, extended modeling was applied to *CL* datasets of 17 drugs, which have been previously modeled using the growth and maturation model of Anderson et al. (24); the model parameters were directly available from the literature (15).The *CL* values were generated using the same general models of Anderson et al. (15, 37), as described in Equation 1, where *P* and *P*_*adult*_ are replaced by *CL* and *CL*_*adult*_. All the drug-specific parameters in the age function (Equation 2) and size function (Equation 3) were taken from the literature (15). For each drug, the *CL* values with random errors were generated for 26 age groups (each of 20 subjects). The procedure to generate the *CL* values with random errors was the same as that described above for *GFR*.

Finally, individual experimental data were retrieved from two pharmacokinetic studies of cefetamet in pediatric patients (47, 48). The first study included 20 infants of 0.19 to 1.44 years (47). The second study included 18 children (3 to 12 years). The adult data (49, 50) were also included to complete the cefetamet dataset.

### 2.4 Model fitting and validation

In this study, the following mixed-power equation was proposed to model the datasets mentioned above.

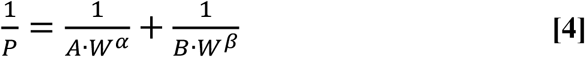

or

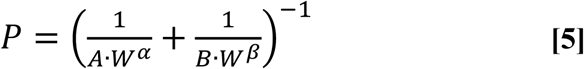

where *P* is the metabolic or physiological quantity of interest, viz. *BMR, GFR* or drug *CL*. The equation consists of two allometric coefficients (0<*A*<*B*) and two allometric exponents (*α*>*β*>0). For model comparison, the simple allometric model was also included for analysis:

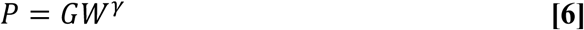

where *G* and *γ* are the allometric coefficient and exponent, respectively.

Model fitting was performed using the nonlinear regression function (*NonlinearModelFit*) in Mathematica^®^ (Wolfram). The model parameters were obtained by fitting the model equations to all available datasets (with suitable weighting functions, e.g., 1/*y*^2^). The comparison of the biphasic allometry (*BA*) model with the simple allometry (*SA*) model was based on the Akaike’s Information Criterion (*AIC*). Besides, the model fitting was validated using multiple measures, including residual plots, *R*^2^, and t-statistic for the fitted parameters.

### 2.5 Estimation of the characteristic values

The biphasic model is characterized by continuous change of slope in the log *P* vs. log *W* plot. The weight-dependent slope (*S*) of the fitted biphasic curve was generated using the following equation (see Appendix for the derivation):

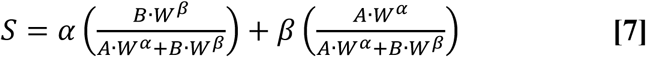

The biphasic growth and maturation curve switches at a characteristic bodyweight (i.e. the bodyweight at the phase transition, 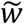), which was calculated using the following expression (Appendix).

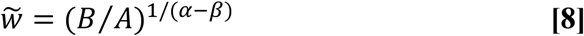

The characteristic slope (*S*\*) at the phase-transition bodyweight was calculated as the sum of the allometric exponents (Appendix), i.e.,

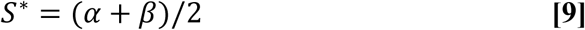

### 2.6 Dimensionless analysis

To conduct a dimensionless analysis, Equation 4 is treated as follows. First, let 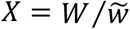, where *W* is the exact bodyweight, and 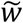 represents the critical bodyweight at the phase transition. Thus, *X* is the dimensionless, fractional number associated with bodyweight. Similarly, let 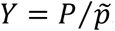, where *P* is physiological parameters such as *BMR* and *GFR*, or *CL*, and 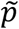 is the corresponding *P* value at 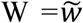, and according to Equation 4,

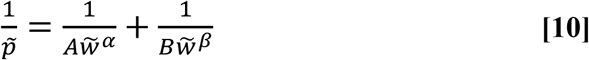

Accordingly, *Y* is defined as the fractional value of *BMR, GFR* or *CL* (relative to their critical value at phase transition). Then, the original Equation 4 can be expressed and rearranged as follows:

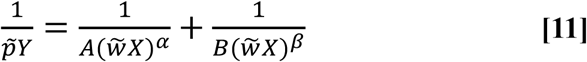

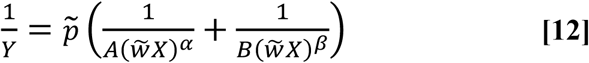

By substituting 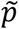 in the last equation from Equation 10, and after further rearrangement, the final equation is obtained:

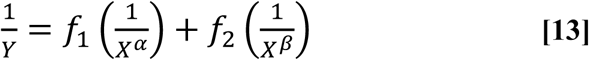

Where

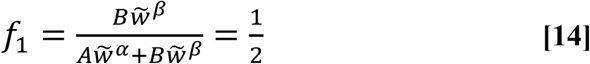

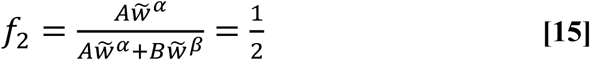

The slope of the dimensionless biphasic-scaling curves (on a log-log plot) has the following form:

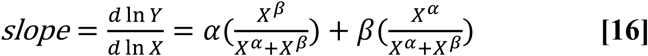

At *X* ≪ 1, the slope is approaching the asymptotic exponent, *α*, whereas at the other end (i.e., *X* ≫ 1) the slope is approaching another exponent, *β*. Since *α* > *β*, the slope continuously decreases as *X* increases. Thus, the decrease of slope (or “deceleration”) is expressed as:

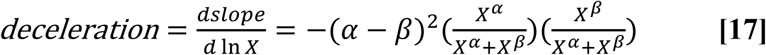

The deceleration equation—which always gives a negative value—quantifies the change (i.e. continuous slowing down) of pace in the growth and maturation process. In this study, different physiological and clearance datasets were harmonized using the derived dimensionless equations and the fitted parameter values for various datasets. Specifically, the dimensionless *Y* values was simulated using Equation 13 with input *X* values ranging from 0.1 to 10 (step of increment = 0.1). The corresponding slope and deceleration curves were generated using Equation 16 and Equation 17, respectively.

## 3. Results

### 3.1 Scaling basal metabolic rate

The biphasic allometry (*BA*) model and the simple allometry (*SA*) model were fitted to the Schofield’s historical dataset of *BMR* (41). Figure 1 shows the fitting results for the male and female datasets. The log *BMR*-*vs*.log *W* plot (Figure 1A and 1B) reveals curvature with two obvious phases. Thus, the mixed-power equation (the biphasic model) offers better fitting results with smaller *AIC* values than the simple allometry equation (Table 1; Figure 1A and 1B). Particularly, the *SA* model systematically overestimates the *BMR* values at *W* <10 kg but underestimates the values at the *W* range of 10 to 40 kg (Figure 1C and 1D). The slope of the biphasic curve decreases with increasing bodyweight; at the extreme body sizes, the slope approaches the two asymptotic allometric exponents, *α* and *β* (Figure 2). The first exponent is about 2, whereas the second exponent is near 0.3 or 0.4 (Table 1). Accordingly, the first phase has a superlinear scaling relationship (slope >1), whereas the second phase scales sublinearly (slope <1). The two phases have a characteristic transition point at 5.5 kg and 6.6 kg, for males and females, respectively (Table 1). Figure 2 highlights the phase transition as the body grows. The slope at the phase transition is the mean of the two asymptotic exponents (Equation 9) (male: 1.22 and female: 1.14).

**Figure 1.**
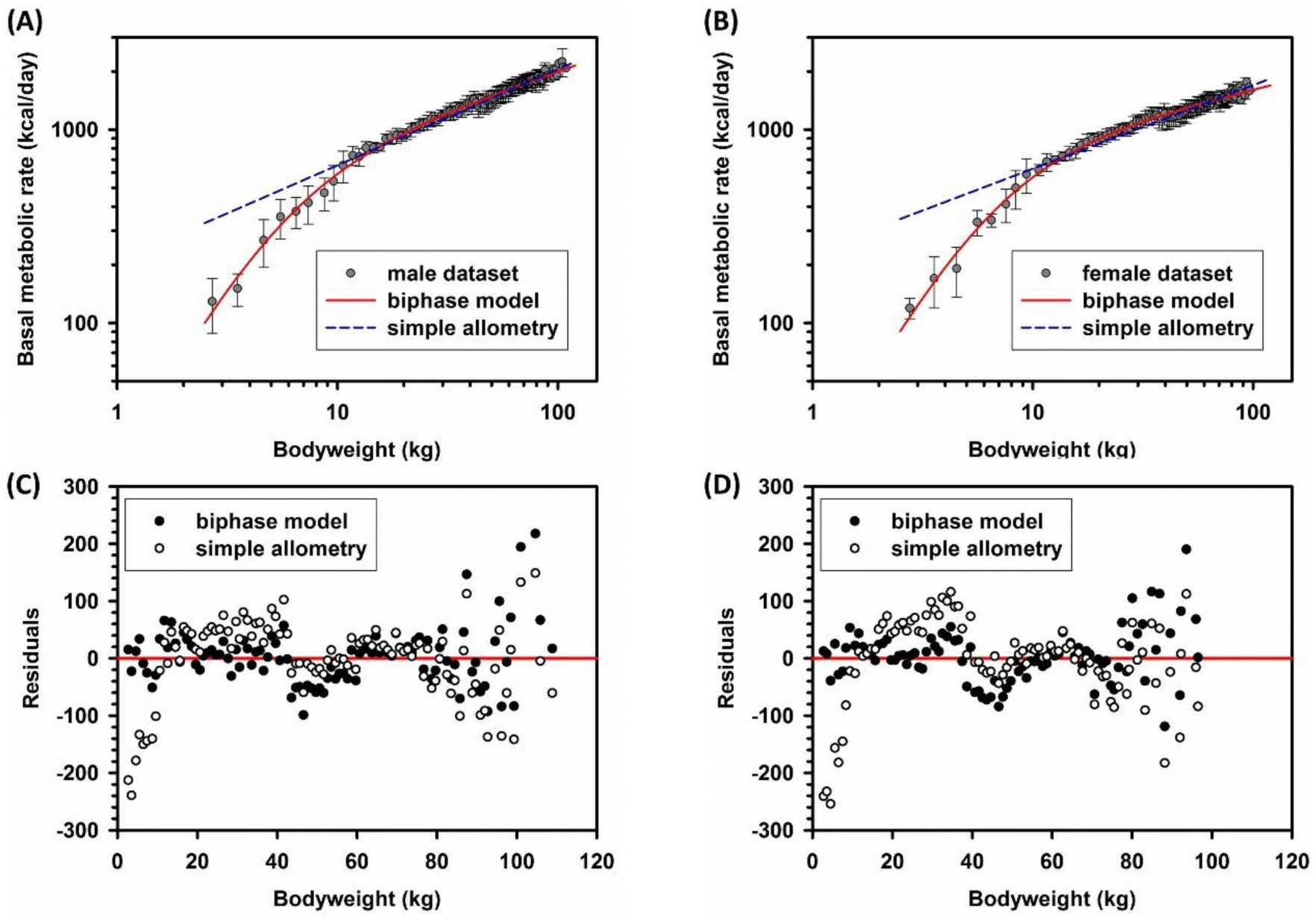
Bodyweight scaling of *BMR* datasets. *BMR* data were retrieved from Schofield’s historical database (41). (A) Comparison of model fittings for the male dataset. Solid line: biphasic model (Equation 4); dashed line: simple allometry model (Equation 6). (B) Comparison of model fittings for the female dataset. (C) The residual plots for the male data. Solid symbol: biphasic model; open symbol: simple allometry. (D) The residual plots for the female data.

**Table 1.** Model fitting results for basal metabolic rate

| | $A$ | $\alpha$ | $B$ | $\beta$ | $\tilde{w}^*$ | $S^*$ | $AIC (BA, SA)$ |
| --- | --- | --- | --- | --- | --- | --- | --- |
| male | 19.5±2.5 | 2.04±0.11 | 327±26 | 0.39±0.02 | 5.52 | 1.22 | (1092, 1151) |
| female | 17.3±2.3 | 2.00±0.10 | 444±44 | 0.28±0.02 | 6.64 | 1.14 | (983, 1063) |

The parameters ( $A, B, \alpha, \beta$ ) are mean ± SE; bodyweight at phase transition, $\tilde{w} = (B/A)^{1/(\alpha-\beta)}$ (Equation 8); the slope at transition, $S^* = (\alpha + \beta)/2$ (Equation 9); Akaike's Information Criterion ( $AIC$ ); biphasic allometry model ( $BA$ ); simple allometry model ( $SA$ ).

**Figure 2.**
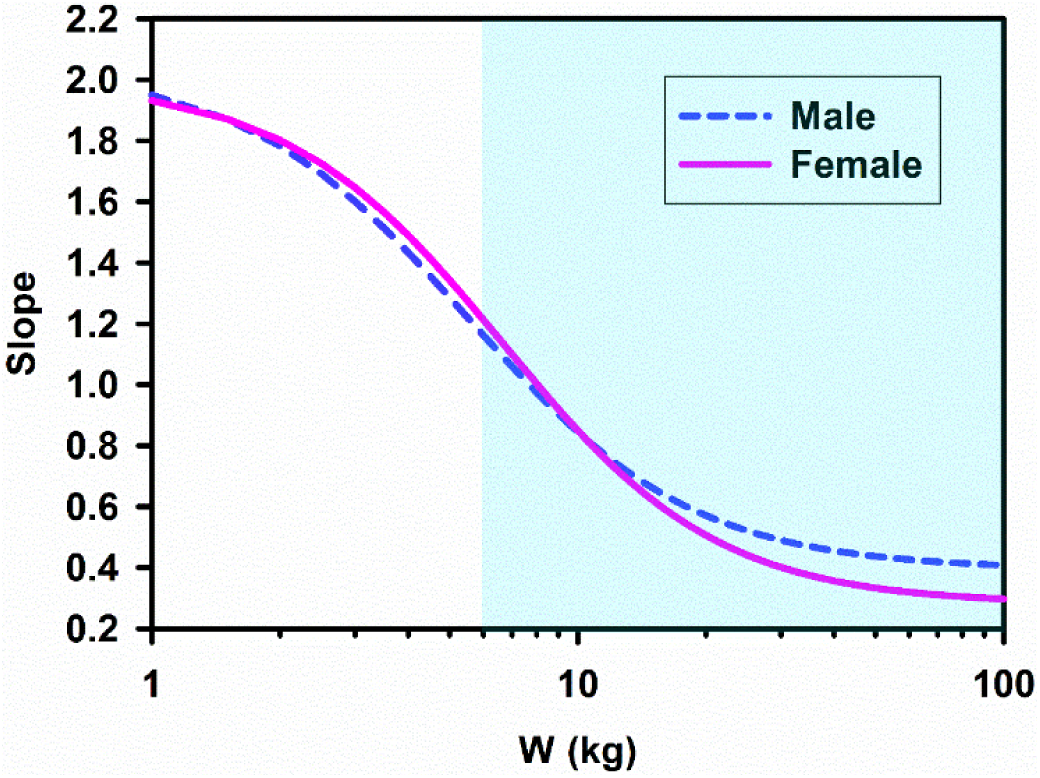
The slope plots for the biphasic model. The changes of the slope on the log-log biphasic plots of the BMR datasets were quantified using Equation 7. The slopes for the males (dashed line) and females (solid line) have a sharp turnaround around 6 kg. The shaded area highlights the transition of the two phases.

### 3.2 Scaling the glomerular filtration rate (*GFR*)

To demonstrate whether the proposed biphasic model could also be applied for scaling *GFR*, the (*GFR, W*) data were simulated for a hypothetical population with 26 age groups. In each group, the bodyweight for 20 subjects was randomly sampled from a lognormal distribution with transformed population mean and estimated standard deviation (Section 2.2). The *GFR* values with random errors (*CV* = 20%) were generated according to Equations 1-3 and using reported parameter values (24), assuming lognormal distribution. The data of all age groups were pooled and then fitted to the biphasic mixed-power equation. Figure 3 shows again a biphasic log-log profiles. The fitted parameter values are: *A* = 0.38 ± 0.07, *α* = 2.18 ± 0.15, *B* = 14.0 ± 3.5, *β* = 0.50 ± 0.06. The result is consistent with a superlinear scaling phase (slope >1) and a sublinear scaling phase (slope <1). The scaling slope decreases with increasing bodyweight (Figure 3A). The phase transition occurs at 8.6 kg and the slope at the transition is 1.34. The simple allometry model overestimates *GFR*, again, at *W* <10 kg (Figure 3B and 3D). Overall, the biphasic model provides better fit with lower *AIC* (3774 vs. 4294) and more randomly scattered residuals (Figure 3C).

**Figure 3.**
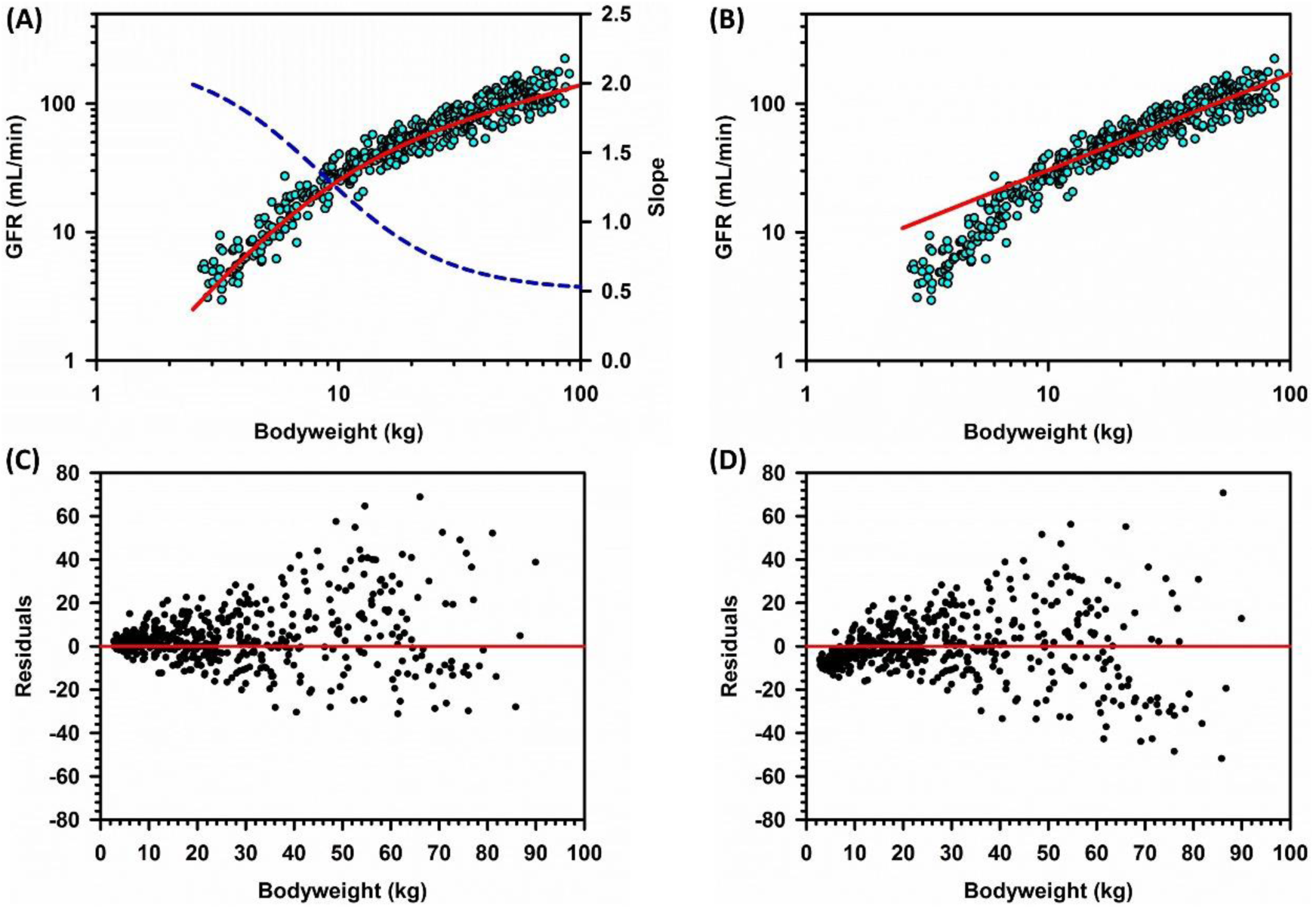
Bodyweight scaling of model simulated *GFR*. *GFR* values with random errors (symbol) for 26 age groups (20 datapoints for each group) were simulated using the growth and maturation model of Andersen et al. (Equations 1-3). (A) The log-log plot for (*GFR, W*) and the slope plot. The solid line represents the nonlinear best-fit for the biphasic model (Equation 4). The dashed line is weight-dependent slope of the fitted curve (Equation 7). (B) The residual plot for the biphasic model. (C) The same data were fitted to the simple allometry model (Equation 6). (D) The residual plot for the simple allometry model.

### 3.3 Scaling drug clearance

Three types of drug clearance data were included for analysis. First, the proposed biphasic model was fitted to the cross-age *CL* data for theophylline (Figure 4 A and 4B) and midazolam (Figure 4C and 4D), which were taken directly from a *PBPK* modeling study (29). The datapoints are scarcely distributed in the bodyweight range spanning neonates and adults. Nevertheless, compared with simple allometry (based on *AIC* and residual plots), the biphasic allometry model provides better fitting results. (Table 2). Again, the features of the biphasic model (i.e. 2 phases with defined phase transition) were clearly illustrated in this analysis. The two drugs have high first-phase exponents (ca. 2.6 to 3) and comparable—but less than unity—second-phase exponents (ca. 0.5 to 0.6). The results also show superlinear allometry in the early childhood and sublinear allometry in the later life. The transition occurs at 8.6 to 9.5 kg, with the corresponding slope values (*S*\*) close to 1.6 and 1.8, for theophylline and midazolam, respectively (Tables 2).

**Figure 4.**
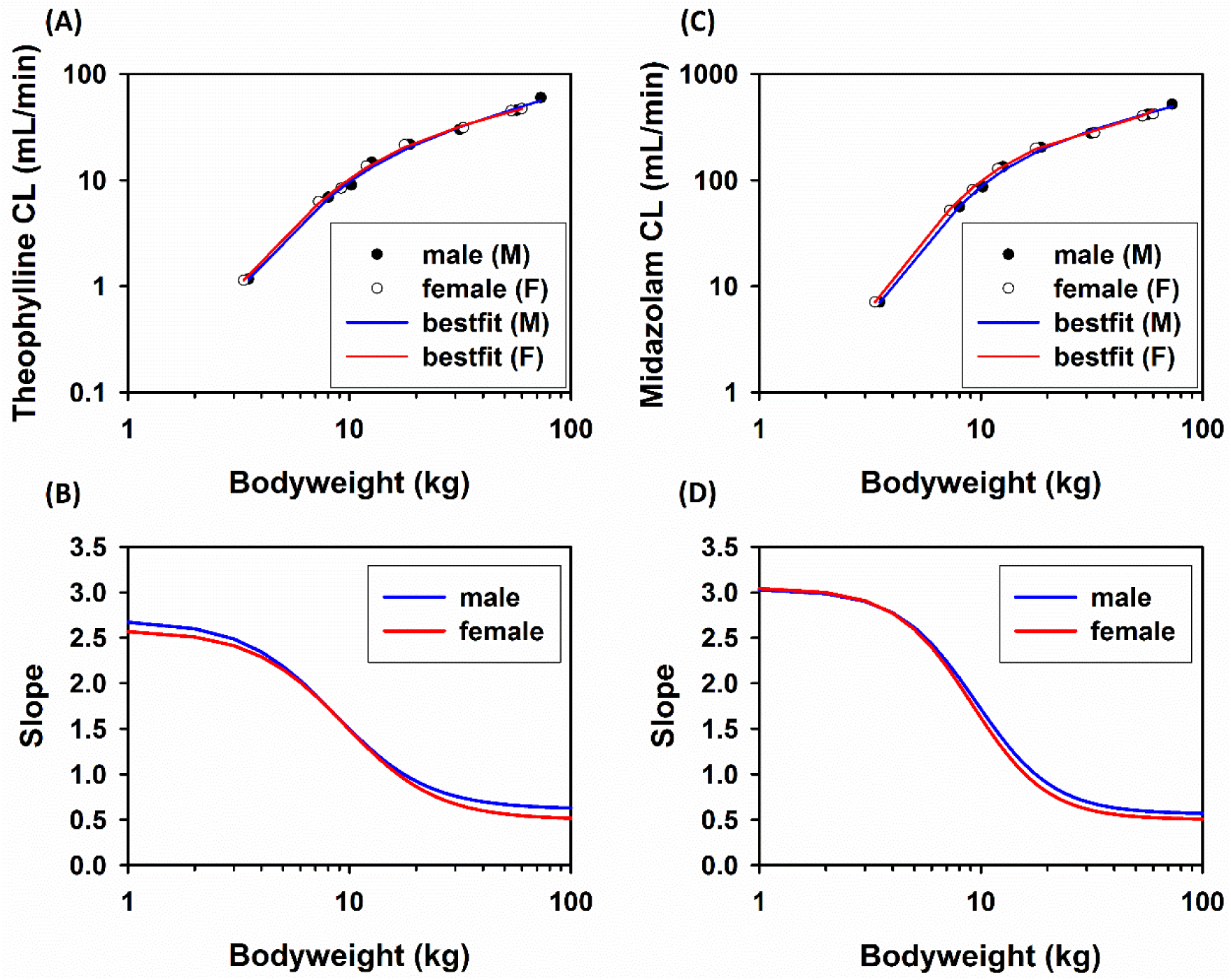
Bodyweight scaling of drug clearance. The log *CL*-vs.-log *W* plots and the slope plots for theophylline (A and B) and midazolam *CL* (C and D). Symbols are *PBPK*-modeled *CL* values directly taken from the literature (29). Lines are the best nonlinear-regression fits for the biphasic model.

**Table 2.** Model fitting results for theophylline and midazolam clearance

| | $A$ | $\alpha$ | $B$ | $\beta$ | $\tilde{w}$ | $S^*$ | $AIC (BA, SA)$ |
| --- | --- | --- | --- | --- | --- | --- | --- |
| Theophylline |  |  |  |  |  |  |  |
| male | 0.05±0.02 | 2.70±0.39 | 4.03±2.69 | 0.62±0.16 | 8.60 | 1.66 | (33.0, 41.5) |
| female | 0.06±0.02 | 2.59±0.22 | 6.18±3.16 | 0.50±0.13 | 9.50 | 1.55 | (25.3, 41.9) |
| Midazolam |  |  |  |  |  |  |  |
| male | 0.17±0.05 | 3.04±0.21 | 44.3±17.1 | 0.56±0.10 | 9.47 | 1.80 | (62.1, 78.5) |
| female | 0.19±0.04 | 3.05±0.14 | 53.8±14.7 | 0.50±0.07 | 9.09 | 1.78 | (55.0, 78.8) |

*PBPK*-modeled CL data were retrieved ref.(29). The parameters ( $A$ , $B$ , $\alpha$ , $\beta$ ) are mean±SE; bodyweight at phase transition, $\tilde{w} = (B/A)^{1/(\alpha-\beta)}$ (Equation 8); the slope at transition, $S^* = (\alpha + \beta)/2$ (Equation 9); Akaike's Information Criterion (*AIC*); biphasic allometry model (*BA*); simple allometry model (*SA*).

Secondly, the analysis was extended to 17 drugs. The *CL* maturation models for these drugs have been previously reported. All the model parameters were retrieved from the literature (15). The maturation and growth model used for generating the *CL* value is independent of the biphasic model proposed in this study. The *CL* values with random errors for 26 age groups (20 subjects/group) were simulated using the same approach described for the *GFR* dataset. For each drug, the population pooled (*CL, W*) data were fitted to both the biphasic model and the simple allometry model.

Table 3 summarizes all fitted parameters. Compared with simple allometry, biphasic allometry fits better, with universally lower *AIC* values (Table 3). The allometric coefficients (*A & B*) are highly variable among drugs, because the absolute *CL* values span >3 orders of magnitude. However, the allometric exponents have a narrow distribution: the first exponent (*α*) ranges from 1.51 to 3.67 (mean = 2.36; *CV* = 26%), whereas the second exponent (*β*) is in the range of 0.40 to 0.67 (mean = 0.55, *CV* = 14%). The mean transitional weight is 10.0 ± 4.9 kg (range = 3.64 to 20.4 kg, *CV* = 49%). The mean transitional slope (*S*\*) is 1.46 ± 0.32 (range = 1.0 to 2.0; *CV* = 22%). The results reveal consistent biphasic bodyweight scaling with superlinear-to-sublinear phase transition. Moreover, although the drugs are highly variable in terms of elimination pathways and the magnitude of *CL* value, the characteristic slope of the second phase (i.e., *β*) is nearly homogenous among different drugs with a CV of 14%.

**Table 3.** Fitted and estimated parameters for drug *CL* datasets

| Drug | $A$ | $\alpha$ | $B$ | $\beta$ | $\tilde{w}$ | $S^*$ | $AIC (BA, SA)$ |
| --- | --- | --- | --- | --- | --- | --- | --- |
| Acetaminophen | 0.980 | 2.05 | 39.7 | 0.50 | 10.8 | 1.28 | (4813, 5317) |
| Acyclovir | 0.042 | 3.67 | 21.8 | 0.63 | 7.88 | 2.15 | (4909, 5330) |
| Busulfan | 1.170 | 1.80 | 24.4 | 0.51 | 10.7 | 1.16 | (4269, 4859) |
| Dexmedetomidine | 0.680 | 2.11 | 45.8 | 0.46 | 12.8 | 1.29 | (4746, 5281) |
| Lamivudine | 0.580 | 2.26 | 30.8 | 0.55 | 10.2 | 1.41 | (4688, 5242) |
| Levetiracetam | 0.256 | 1.91 | 6.05 | 0.57 | 10.5 | 1.24 | (2991, 3530) |
| Levobupivacaine | 8.318 | 1.71 | 49.6 | 0.59 | 4.93 | 1.15 | (5511, 5970) |
| Midazolam | 0.238 | 2.59 | 52.9 | 0.48 | 12.9 | 1.54 | (5013, 5597) |
| Morphine | 1.732 | 2.57 | 125 | 0.58 | 8.61 | 1.58 | (6333, 6906) |
| Pantoprazole | 1.383 | 1.51 | 31.0 | 0.48 | 20.4 | 1.00 | (4079, 4624) |
| Penciclovir | 0.045 | 3.10 | 63.6 | 0.47 | 15.9 | 1.79 | (5134, 5814) |
| Propofol | 20.13 | 1.84 | 129 | 0.61 | 4.59 | 1.23 | (6598, 7052) |
| Pyrimethamine | 0.019 | 3.31 | 1.23 | 0.67 | 4.90 | 1.99 | (2063, 2586) |
| Ropivacaine | 0.084 | 3.11 | 40.9 | 0.53 | 11.0 | 1.82 | (5070, 5541) |
| Sulfadoxine | 0.005 | 2.11 | 0.04 | 0.67 | 3.70 | 1.39 | (-1720, -1279) |
| Valdecoxib | 0.037 | 2.63 | 20.3 | 0.40 | 16.8 | 1.52 | (3574, 4279) |
| Vancomycin | 1.117 | 1.92 | 5.87 | 0.63 | 3.64 | 1.28 | (3433, 3916) |
| Mean |  | 2.36 |  | 0.55 | 10.0 | 1.46 |  |
| SD |  | 0.63 |  | 0.08 | 4.9 | 0.32 |  |
| CV (%) |  | 26.4 |  | 14.4 | 49.0 | 21.9 |  |

$\tilde{w} = (B/A)^{1/(\alpha-\beta)}$ (Equation 8); $S^* = (\alpha + \beta)/2$ (Equation 9)

Since the bodyweight at the phase transition has a wider distribution, the 17 drugs were further divided into three groups (Figures 5-7). The bodyweight at the phase transition is around 15 kg for the group-I drugs, 10 kg for the group-II drugs, and <5 kg for the group III drugs.

**Figure 5.**
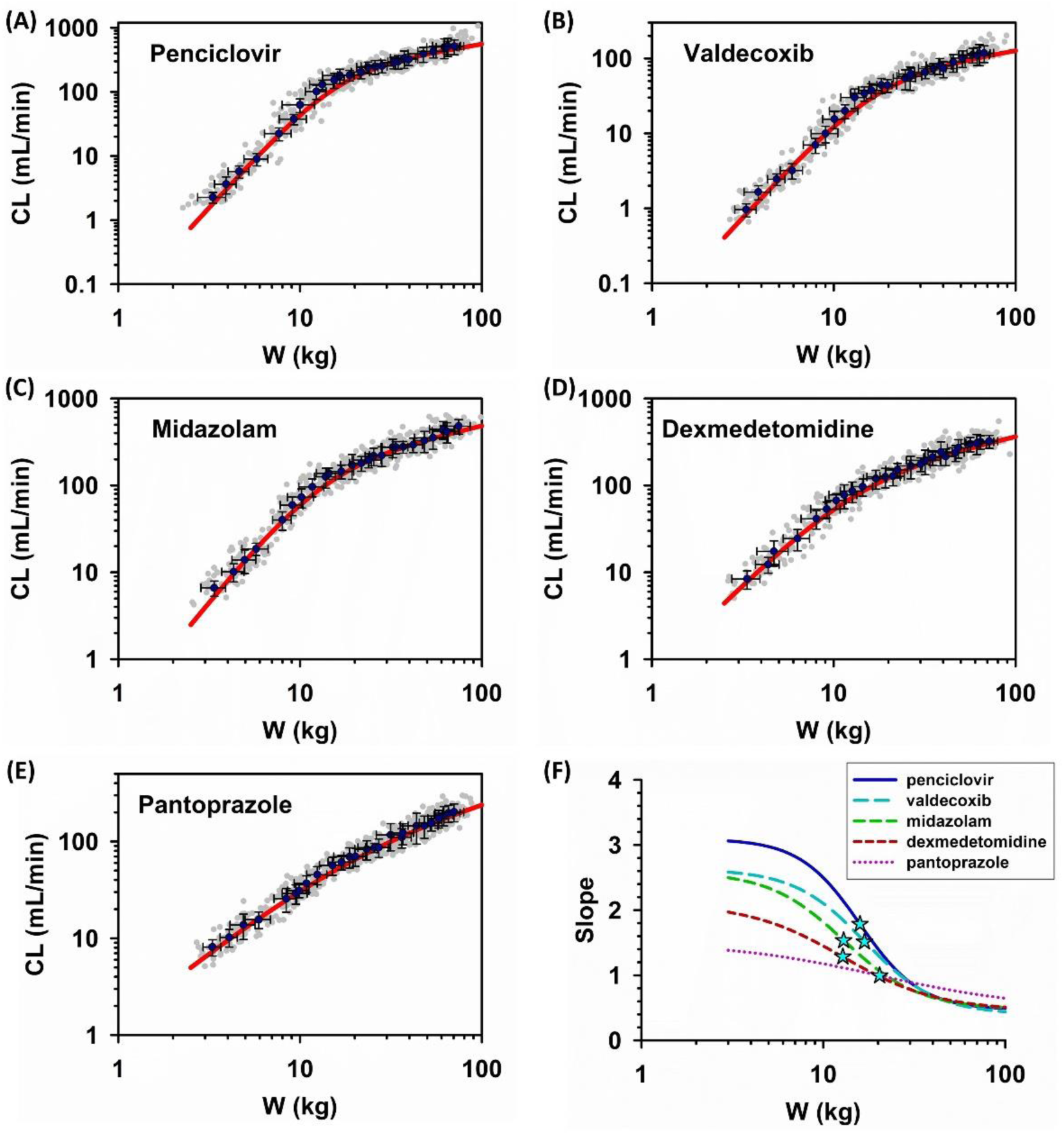
Biphasic bodyweight scaling of *CL* (group-I drugs). Group-I drugs are the drugs with the largest transitional bodyweight (around 15 kg). (A-E) The log *CL* vs. log *W* plots (symbol) and the best-fit lines of the biphasic model. *CL* values with random errors (*CV* = 20%) were generated according to the sigmoidal maturation model (Equations 1-3). Each dataset was fitted to the biphasic model. (F) The slope plots (lines) and the transitional bodyweights (stars) for Group-I drugs.

**Figure 6.**
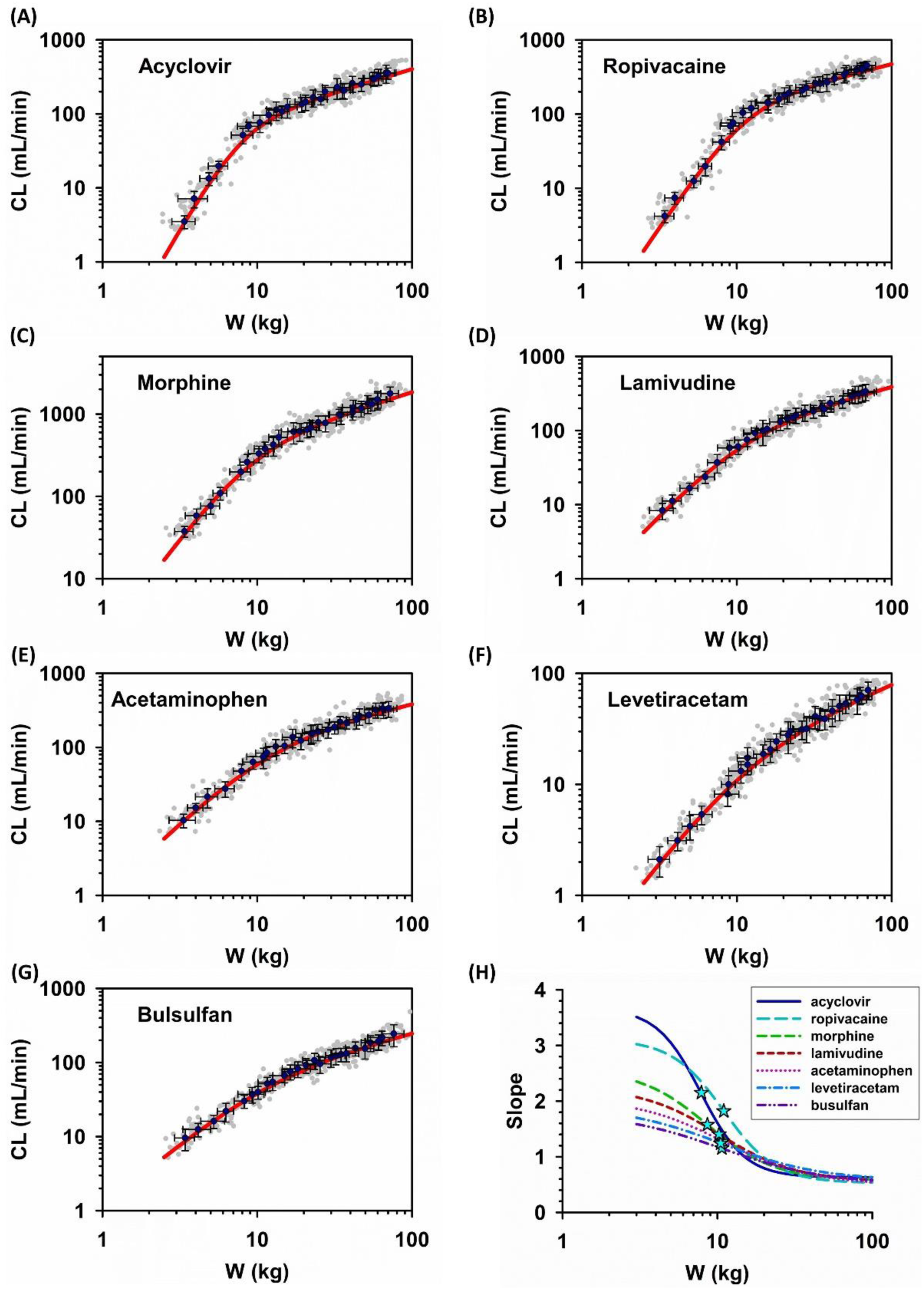
Biphasic bodyweight scaling of *CL* (group-II drugs). Group-II drugs are the drugs with intermediate transitional bodyweight (around 10 kg). (A-E) The log *CL* vs. log *W* plots (symbol) and the best-fit lines of the biphasic model. (F) The slope plots (lines) and the transitional bodyweights (stars) for Group-II drugs.

**Figure 7.**
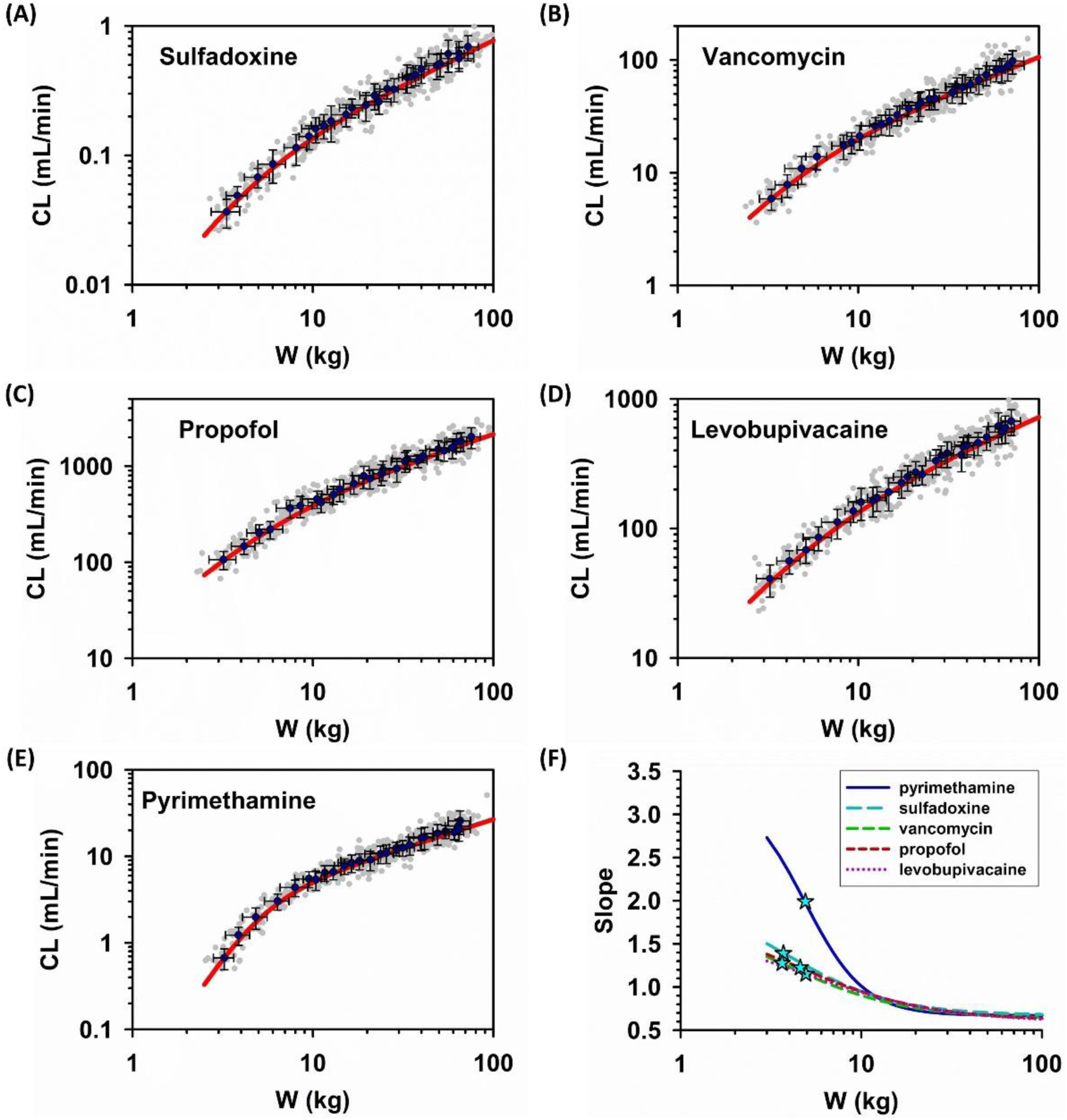
Biphasic bodyweight scaling of *CL* (group-III drugs). Group-III drugs are the drugs with the lowest transitional bodyweight (<5 kg). (A-E) The log *CL* vs. log *W* plots (symbol) and the best-fit lines of the biphasic model. (F) The slope plots (lines) and the transitional bodyweights (stars) for Group-III drugs.

Finally, to demonstrate the versatility of the biphasic model, the drug clearance values from individual pediatric subjects were retrieved from two studies of cefetamet (47, 48). Figure 8 shows that biphasic model fits all the datapoints well, except for one point (which is from a subject that had multiple postoperative complications (48)). The fitting results are described in the figure legend. While *A* and *B* values are drug specific, the *alpha* exponent (2.9) falls within the range of the values found for other drugs (Table 3). Moreover, the beta exponent (0.47) and the transitional bodyweight (11 kg) are about the mean values estimated for the datasets of 17 drugs.

**Figure 8.**
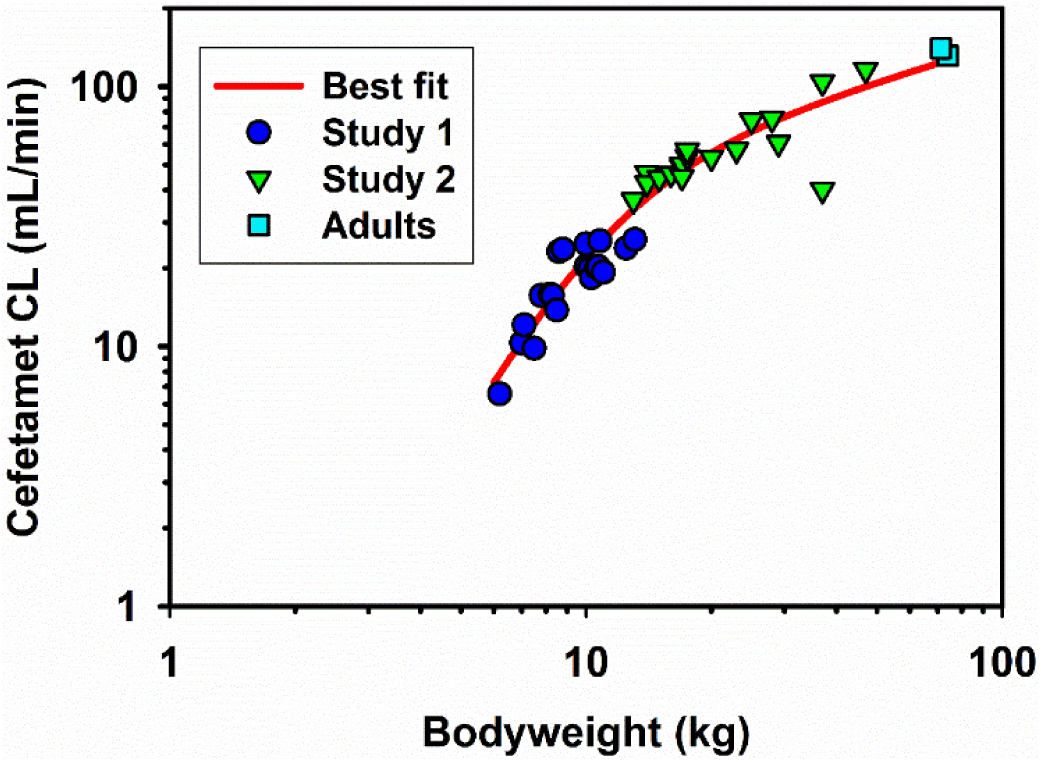
Biphasic bodyweight scaling of individual pediatric *CL* data obtained from different pharmacokinetic studies. The individual *CL* values of cefetamet were retrieved from study-1 (20 infants, 0.19 to 1.44 years)(47) and study-2 (18 children, 3 to 12 years)(48). The adult data were from ref. (49, 50). One subject in the second study was reported to have multiple postoperative complications (coincidentally, the one that has a *CL* value far below the fitted line). The pooled data were fitted with the biphasic allometry and the simple allometry model. The *AIC* values are 269 (biphasic) vs. 313 (simple allometry). The fitted parameter values for the basic model are: *A* = 0.049, *α* = 2.90, *B* = 16.6, *β* =0.47. The estimated transition bodyweight is 11.0 kg.

### 3.4 Unifying all the datasets

In this study, a total of 25 datasets were included for analysis. These data cover 3 types of human parameters (*BMR, GFR* and drug *CL*), where the absolute values span >3 orders of magnitude. Nevertheless, some common features repeatedly appear throughout this study. To further highlight the similarities and differences among different datasets, a dimensionless analysis was performed, according to the definition of *X* and *Y*, and the derived equations (Equations 13, 16, and 17; Section 2.6). The two new variables (*X, Y*) are fractions or multiples of the two characteristic values (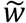 and 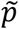). With the dimensionless equation (e.g., Equation 13), all of the 25 datasets can be compared in one figure (Figure 9A). This composite plot clearly divides the biphasic curves into two segments at the reference point of (*X, Y*) = (1, 1), which is considered as a phase transition (turnaround) point. The first segment is within the region where both *X* and *Y* are below the unity, whereas the second segment locates at (*X, Y*) above unity. All the data in the first phase show steeper slopes than those in the second phase (i.e. *α* > *β*). Moreover, the first phase is more variable among different data than the second phase. This is consistent with the magnitude of variability reported in Table 3 for the drug *CL* (*CV* for *α* = 26.4%; for *β* = 14.4%).

**Figure 9.**
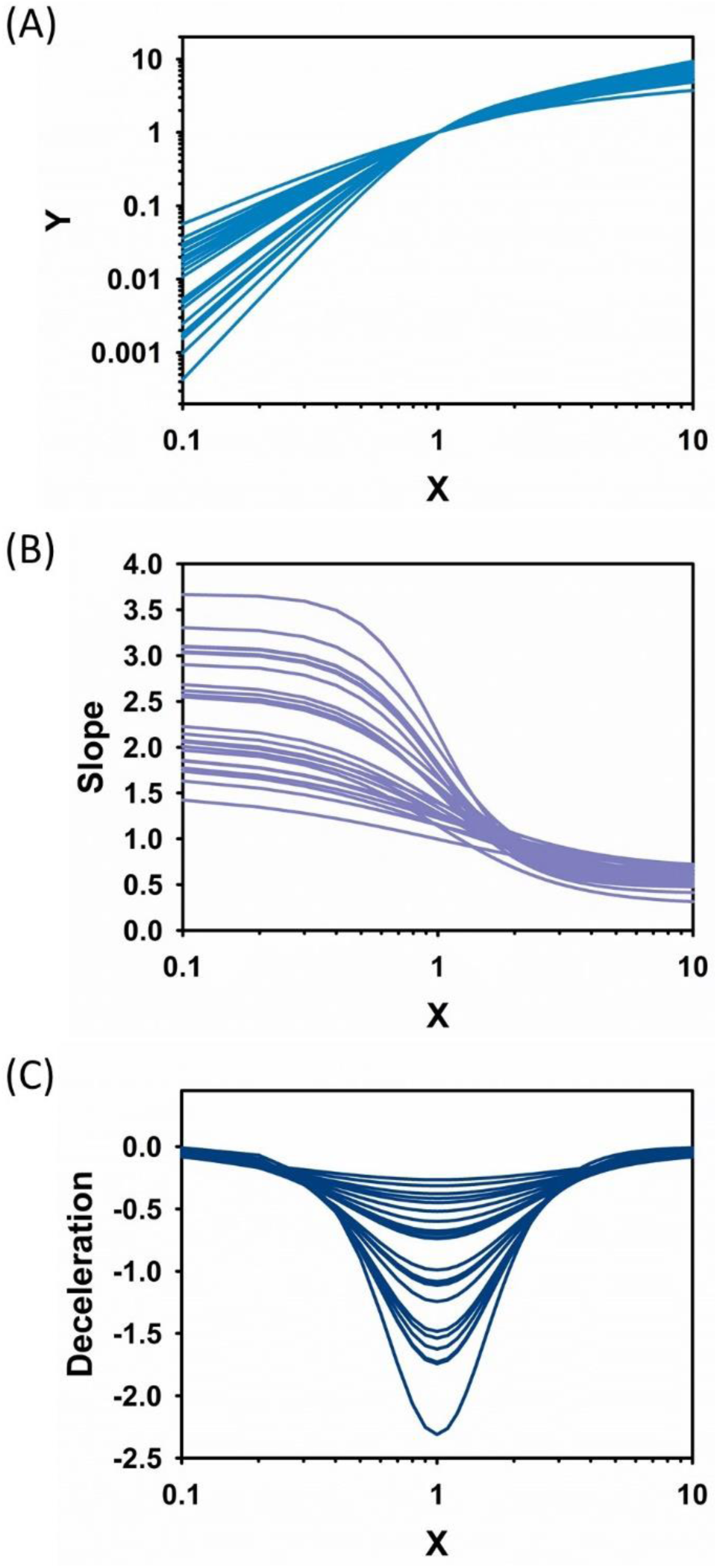
Unifying 25 datasets using dimensionless analysis. (A) The log-log plot for the dimensionless variables. (B) The slope plots. (C) The deceleration plots. The definitions of the dimensionless variables, *X* (fractional *W*) and *Y* (fractional *P*), and of the slope and deceleration are described in Section 2.6. For each individual dataset, the fitted parameters of the biphasic model were used to simulate the dimensionless data, using Equations 13, 16, and 17. A total of 25 datasets (covering *BMR, GFR* and drug *CL*) were compared in one plot.

Figure 9B further captures the essence of such disparity; while the asymptotic slope in the first phase is distributed from 1.5 to 3.5, the asymptotic slope in the second phase has a narrow distribution (ca. 0.4 to 0.7). A consistent observation is that all the data seemed to merge at *X* >2 (i.e., *W* is 2 times that of the transitional bodyweight), suggesting that the maturation and growth patterns of various human parameters are similar when the growing body is passing the critical point of transition.

Figure 9C offers further insights into the deceleration process of human development after birth. By definition (Section 2.6), deceleration is the change of the slope of the maturation curve. The negative value suggests that, although the human function is growing, the speed of functional maturation is continuously declining, i.e. deceleration. Figure 9C shows many characteristics of the deceleration process. First, it is an inverse, symmetrical bell-shaped curve with maximal deceleration occurring at the critical turnaround point (1, 1). Second, slow deceleration in the early life when the body size is small, and when the adulthood is approaching. Third, fast deceleration starts almost universally at *X* = 0.3 to 0.4, reaching the maximal deceleration at *X* = 1. This characterizes the maturation pattern of the first phase. After passing the maximal deceleration point, the maturation enters the second phase, and again, approaches zero deceleration at the adulthood (completing the maturation process).

## 4. Discussion

The current study proposes a general weight-only equation to model various ontogeny datasets of human parameters. The datasets are diverse, spanning multiple human parameters and covering extensive age groups from birth to adulthood. The data were either directly retrieved from historical datasets (including mean and individual values) or simulated from independent models with published values for specific model parameters. A statistical random sampling approach was used to add random errors to the simulated data. The results demonstrate that a biphasic, mixed-power-law equation with two limiting allometric functions adequately quantifies the developmental phase-transition profiles of *BMR, GFR* and drug *CL*. The biphasic model and the additional dimensionless analysis offer quantitative insights into the maturation and growth of human body functions.

The long-held belief that body functions develop at distinct pace in different stages of human development is well supported by extensive empirical evidence (1, 5, 51-53). For example, a study dated back to 1949 had shown that renal function matures with the most rapid rate in the first 6 months, followed by slower rates to reach adult values around the second year of life (51). Numerous empirical models have been proposed to scale the maturation and growth processes of body functions (6, 7, 24). These quantitative equations have been frequently developed for practical considerations, e.g., estimating children’s values (6, 14, 39). The equations are usually data-specific and age-categorized. Most equations contain bodyweight as an input variable; meanwhile, age was often included as an additional variable (6-8, 24, 36). The equations are either separate, discontinuous equations with cut-points at certain age thresholds (23, 40), or continuous nonlinear equations containing age and bodyweight terms (7, 8, 15, 24). The dependence of bodyweight on age (54, 55) often required to assume a fixed weight exponent value (i.e. 0.75) in order to estimate the age-associated parameters (e.g., maturation half-life and Hill coefficient) (15, 24). Accordingly, this present study provides a different perspective where bodyweight alone in a continuous 4-parameter power-law equation is sufficient to quantitatively describe the maturation process.

A previous study has shown that brain’s metabolic requirement starts to accelerate after the first 6-month of life and reaches a lifetime peak at about 5 years of age (1). During this particular period, the increasing demand of energy in human brain development was compensated by slow rate of body growth (1). This present study reveals that the rate switching of *BMR, GFR* and *CL* occurs at bodyweights of around 5-20 kg, which is about the bodyweight range for children of 0.5-9 years of age. Moreover, the transitional bodyweights for different datasets center around 10 kg, which is about the weight of children at 2 years of age.

The biphasic profiles are universally reproduced in different datasets. Since other important metabolic organs, such as the liver and the kidneys, may develop similar age-related dynamics in energy demands, the biphasic profiles may be associated with the trade-off between maturation (functional development) and growth (body-size development). In the first developmental phase, high blood-perfusion organs (e.g., brain, liver, kidney) take up more metabolic substrate (e.g., glucose) to allow fast development of organ functions (fast maturation), which results in less energy available for the building of body mass (slow growth). Therefore, this phase is characterized by high energy needs for the maturation of vital organs, leading to the superlinear bodyweight scaling relations (exponent >1). However, as the body size grows, the energy supply relative to the maturing vital organs might need to be bounded or be re-shuffled to other organs e.g. muscle. This may contribute to continuous decrease of metabolism and many other organ functions along the maturation process (56). This is considered as a deceleration process in this study. The metabolic deceleration continues through the second phase but with a pace different from that of the first phase. Together, energy re-allocation between high and low energy-demanding organs may attribute to biphasic maturation and growth of organ functions. This view is consistent with the general model of West et al. (54) which suggests that the growth of body size is limited by the capacity of resource allocation networks.

As illustrated in Appendix (Figure A1), the curvature of the biphasic profile is set by the two allometric functions. Each of the two extreme states is represented by a single allometric function of bodyweight raised to a constant power (*α* or *β*). The magnitude of the two exponents shape the entire curve: for example, higher *α* together with lower *β* produces greater curvature and more obvious phase transitions, as indicated in the modeling of the *CL* values for 17 drugs (Table 3 and Figures 5-7). Moreover, the biphasic model shows that the slope of the scaling curve continuously changes during the developmental period, which agrees with previous observations (16, 28, 30).

The essence of the phase switching with deceleration is further captured by the dimensionless analysis. According to Equation 13, all unitless data will converge at (*X, Y*) = (1, 1), which is also the transition point of the biphasic curve. The dimensionless equations indicate that metabolic and physiological functions scale with bodyweight superlinearly at *X*<1 (i.e. bodyweight < transitional weight) and sublinearly at *X*>1 (i.e. bodyweight > transitional weight). The dimensionless analysis can be further elaborated as follows. Mathematically, Equation 16 can be expressed as the linear combination of two fractional terms:

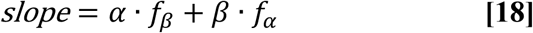

where

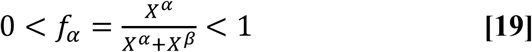

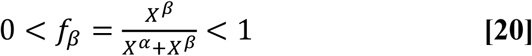

and the deceleration equation (Equation 17) can be simplified as:

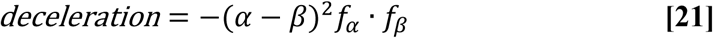

In biological terms, the two fractions represent the relative contribution of the two maturation/growth phases (i.e. *alpha* and *beta* phases) from birth to adulthood. Phase transition occurs at the characteristic bodyweight where the two phases have equal contribution (i.e. *f* = 1/2). Thus, at the phase transition point, the slope is the average of the two characteristic exponents, i.e., (*α* + *β*)/2, and deceleration reaches the maximal value: −0.25(*α* − *β*)^2^. In this study, the *beta* exponent is around 0.5 with small variation across all 25 datasets. In contrast, the *alpha* exponent has larger variation (Table 3 and Figure 9B). Thus, larger difference in the two exponents results in sharper transition and greater peak deceleration (Figure 9C).

Finally, some caveats and future perspectives are noted here. The aim of this study is *<u>not</u>* to predict the unknown values from the proposed model. Instead, the study is the first attempt aiming to understand the ontogeny process of human metabolism and physiological functions by proposing a general, harmonizing model. Future studies may be directed towards building and validating the proposed model for practical applications, such as prediction of renal functions, of pediatric doses and of metabolic demands. Such studies would have broad implications in drug therapy and clinical nutrition. Moreover, the proposed biphasic model may be applied in future studies of population pharmacokinetics and PBPK modeling. Finally, this paper may inspire future attempts in finding a theoretical framework and derivation of the equation based on first principles.

## 5. Conclusion

In this paper, a general equation is proposed to model the maturation of *BMR, GFR* and drug *CL*. The equation (Equation 4) is expressed in reciprocal terms of mixed allometric functions where bodyweight is the only independent variable. All the bodyweight-based profiles are biphasic on log-log coordinates. The biphasic curvature is determined by two asymptotic allometric functions, representing the maturation and growth functions, respectively. The first phase (below the critical bodyweight) is universally characterized by much steeper slopes (i.e. superlinear scaling with slope >1). The second phase (above the critical bodyweight), however, exhibits sublinear scaling (slope <1). The proposed equation can be further expressed in a unitless form, where the relationship between the fractional physiological/metabolic quantity (*Y*) and the fractional bodyweight (*X*) can be described by a general equation with two allometric exponents, *α* and *β*. The unified equation quantitatively characterizes the curvature of the biphasic maturation and growth profiles for diverse datasets, by two characteristic parameters (*α* and *β*). In conclusion, the proposed mixed-allometry model enables a quantitative understanding of human development from birth to adulthood.

## Acknowledgments

The author is supported by Ministry of Science and Technology (MOST) and Higher Education Sprout Project – Ministry of Education (MOE), Taiwan.

There is no competing interest to declare.

## 7. Appendix

### Characteristics of the biphasic model

This study proposes a mixed-power function (the biphasic allometry model; Equation 4) to scale human *BMR, GFR* and drug *CL* from adults to neonates. In fact, this study was inspired by allometric scaling of whole-plant respiration from small seedlings to giant trees (12). Equation 4 states that the reciprocal of a physiological or metabolic parameter (*P*) is equal to the sum of reciprocals of each of two individual power-law bodyweight functions (i.e. allometric functions). To understand the key features of the model, a simulation was conducted to characterize the mixed-power function. The following hypothetical parameters were used: *A* = 0.1, *B* = 10, *α*= 2.5, *β*= 0.5. By plugging the parameter values into Equation 4, a set of hypothetical (*P, W*) data was generated for a simulated body-weight range of 3-70 kg (at a 1-kg interval). Figure A1 (A) shows the simulated data and two asymptotic lines on log-log coordinates. Let *P*_1_ and *P*_2_ be the first and second asymptotic functions, respectively, then the asymptotic bodyweight functions are:

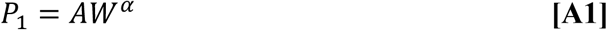

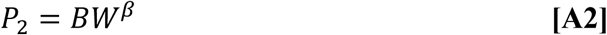

and Equation 4 can be expressed as:

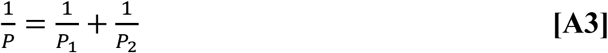

When *P*_1_ ≪ *P*_2_, *P* ≈ *P*_1_, which is the first asymptote near the low bodyweight region of *W* ≪ (*B*/*A*)^1/(*α*−*β*)^ (by solving the inequality relationship). In contrast, when *P*_1_ ≫ *P*_2_, *P* ≈ *P*_2_, and the second asymptote occurs at *W* ≫ (*B*/*A*)^1/(*α*−*β*)^. Thus, the characteristic bodyweight 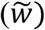 is revealed as:

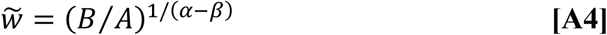

Mathematically, 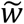 is exactly the bodyweight at which the two asymptotic lines intersect (*P*_1_ = *P*_2_ and 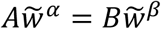). Biologically, it is considered here as the critical bodyweight at which the body undergoes transition from maturation phase to growth phase (i.e. the phase-transition bodyweight). By taking the derivative of Equation 4, the following expression for the slope (*S*) of the biphasic curve is obtained:

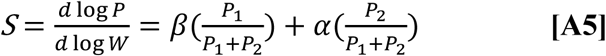

Therefore, according to Equation A5, the slope of the biphasic model can be estimated at any given bodyweight. Figure A1 (B) shows that the slope decreases with increasing bodyweight. The slope curve is indeed flanked by two asymptotic lines: slope ≈ *α*, at the low body weight region where 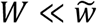 or when *P*_1_ ≪ *P*_2_ ; whereas slope ≈ *β*, at 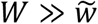 or when *P*_1_ ≫ *P*_2_. Note that the characteristic slope (*S*\*) at the phase-transition bodyweight is exactly the mean of the two asymptotic exponents, since at 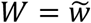, *P*_1_ = *P*_2_, Equation A5 is reduced to:

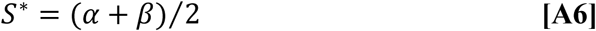

In sum, Figure A1 captures the essence of the proposed mixed-allometry model by revealing all the parameter values used for the simulation: two asymptotic exponents of 2.5 and 0.5 (exactly *α* and *β*), two asymptotic allometric coefficients (*A* = 0.1, *B* = 10), a critical bodyweight of 10 kg (i.e. 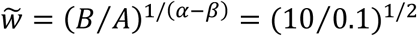) where the slope of the biphasic curve at this critical point is exactly the mean of the two exponents (i.e. 1.5).

**Figure A1.**
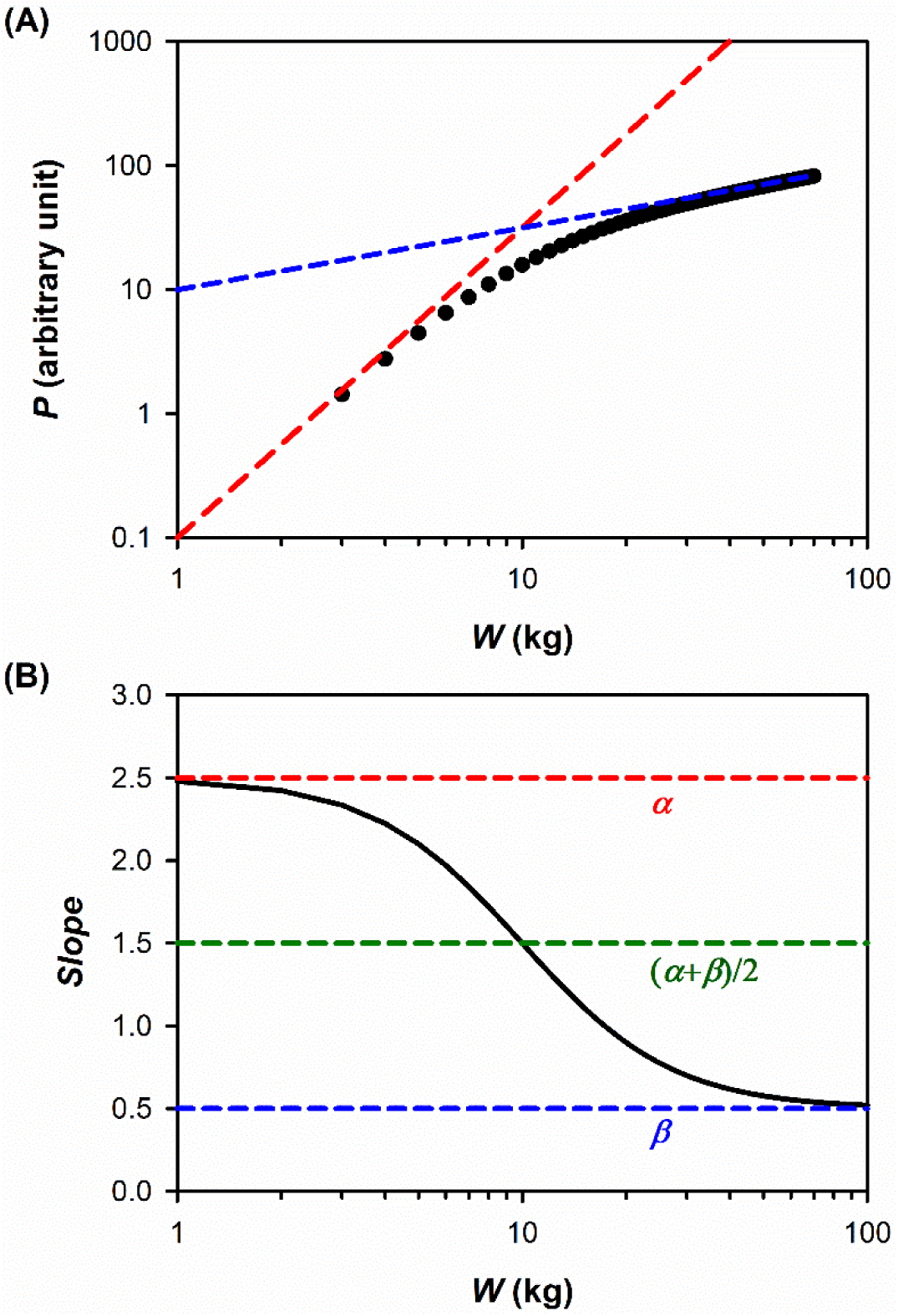
Characteristics of the mixed-power function. (A) Simulated data of physiological quantity (*P*) vs. bodyweight (*W*). The lines represent the two asymptotic power-law functions with respective y-intercepts (i.e., 0.1 and 10, at *W* = 1 kg) and an interception point which defines the critical point of phase transition (at *W* = 10 kg). (B) Slope as a function of bodyweight. The plot highlights the two allometric exponents (*α* and *β*) and the corresponding mean value at the point of phase transition.

